# Modelling and rescue of RP2 Retinitis Pigmentosa using iPSC Derived Retinal Organoids

**DOI:** 10.1101/2020.01.28.923227

**Authors:** Amelia Lane, Katarina Jovanovic, Ciara Shortall, Daniele Ottaviani, Anna Brugulat Panes, Nele Schwarz, Rosellina Guarascio, Matthew J. Hayes, Arpad Palfi, Naomi Chadderton, G. Jane Farrar, Alison J. Hardcastle, Michael E. Cheetham

## Abstract

RP2 mutations cause a severe form of X-linked retinitis pigmentosa (XLRP). The mechanism of RP2 associated retinal degeneration in humans is unclear, and animal models of RP2 XLRP do not recapitulate this severe phenotype. Here, we developed gene edited isogenic RP2 knock-out (RP2 KO) induced pluripotent stem cells (iPSC) and RP2 patient derived iPSC to produce 3D retinal organoids as a human retinal disease model. Strikingly, the RP2 KO and RP2 patient derived organoids showed a peak in rod photoreceptor cell death at day 150 (D150) with subsequent thinning of the organoid outer nuclear layer (ONL) by D180 of culture. AAV mediated gene augmentation with human RP2 rescued the degeneration phenotype of the RP2 KO organoids, to prevent ONL thinning and restore rhodopsin expression. Notably, these data show that 3D retinal organoids can be used to model photoreceptor degeneration and test potential therapies to prevent photoreceptor cell death.

## Introduction

The reprogramming of patient derived cells into induced pluripotent stem cells (iPSC) has enabled the derivation and differentiation of a range of somatic cell types and has revolutionized our ability to study inherited disease (Takahashi et al., 2007). The differentiation of iPSC towards retinal lineages has seen huge advances in recent years with the refinement of protocols for the generation of three dimensional retinal organoids (Nakano et al., 2012, Gagliardi et al., 2019). Unlike previous models in 2D, these 3D structures contain photoreceptors with morphologically identifiable features; including, inner segments rich in mitochondria, rudimentary outer segments with connecting cilia, and synaptic pedicles in addition to bipolar, Müller glia, ganglion and amacrine cells, synaptic layers and an outer limiting membrane, arranged in retinal layers (reviewed in (Capowski et al., 2019). These advanced models have proven to have many translational research applications, including transplantation studies (Shirai et al., 2016, Gonzalez-Cordero et al., 2017), retinal disease modelling and testing efficacy of potential therapies in human photoreceptor cells (Parfitt et al., 2016, Schwarz et al., 2017, Sharma et al., 2017, Deng et al., 2018). To date, however, they have not been used to model and rescue photoreceptor cell death.

Mutations in *RP2* account for approximately 15% of all cases of X-linked retinitis pigmentosa (XLRP) (Hardcastle et al., 1999, Breuer et al., 2002). RP2 is a GTPase activating protein (GAP) for the small GTPase ARL3 (Veltel et al., 2008), which is also regulated by its guanine nucleotide exchange factor (GEF) ARL13B (Gotthardt et al., 2015). ARL13B is localized to the ciliary axoneme, whereas a pool of RP2 and ARL3 localise at the basal body and associated centriole at the base of photoreceptors (Grayson et al., 2002, Evans et al., 2010). ARL3, with its effectors (UNC119 and PDEdelta (PRBP)) and GAP RP2, are thought to be important in the retina to traffic lipidated proteins, such as transducin, GRK1 and PDE6, to the photoreceptor outer segment (Ismail et al., 2011, Wright et al., 2011, Zhang et al., 2011, Schwarz et al., 2012, Zhang et al., 2015). RP2 knockout mice have a relatively mild phenotype compared to human disease. In one model mis-localization/absence of GRK1 and cone PDE6a was evident at 14 months (Zhang et al., 2015), whereas another model was reported to have rhodopsin and M opsin mis-localization at 2 months and ONL thinning at 5 months (Li et al., 2013). In contrast, the human phenotype is relatively severe with some patients experiencing macular atrophy in childhood (Jayasundera et al., 2010), highlighting the necessity for human retinal models of disease.

Currently there are no treatments for this condition, so there is a need to develop potential therapies. Characterization of iPSC derived RPE and early stage retinal organoids from an individual carrying a nonsense mutation in *RP2* (c.358C>T, p.R120X) showed changes in Golgi cohesion, Gbeta trafficking in RPE and ciliary trafficking of Kif7 in retinal organoids (Schwarz et al., 2015, Schwarz et al., 2017). Furthermore, treatment with the readthrough drugs, G418 and/or Ataluren (PTC124), could restore detectable full length RP2 protein and rescue the Golgi cohesion and Gbeta mis-localization in iPSC-RPE and kinesin traffic in retinal organoids. Gene therapy for other inherited retinal diseases using adeno associated viruses (AAVs) have been shown to efficiently transduce photoreceptors and RPE following or subretinal injection (Sarra et al., 2002) in animal models. There is an FDA and EMA approved AAV mediated ocular gene therapy (Russell et al., 2017) and a number of AAV mediated ocular gene therapies are currently in phase I/II and III clinical trials (clinicaltrials.gov). AAV delivery of human RP2 to a mouse knock-out model of RP2-XLRP preserved cone function, but had no effect on rod cell function and toxicity was observed at a higher viral dose (Mookherjee et al., 2015).

Here, we describe the temporal maturation of CRISPR gene edited *RP2* knockout retinal organoids relative to their isogenic control, in addition to retinal organoids derived from two unrelated individuals with the same R120X nonsense mutation. These studies reveal that the loss of RP2 leads to rod photoreceptor degeneration that can be rescued by AAV delivery of RP2.

## Results

### *RP2* knock out and *RP2* patient iPSCs develop mature retinal organoids

Fibroblasts from two unrelated individuals (R120X-A and R120X-B) carrying the nonsense mutation c.358C>T; p.R120X were reprogrammed into iPSC by nucleofection (Okita et al., 2011, Schwarz et al., 2015). CRISPR–Cas9 with guides designed to target exon 2 of *RP2* were used to generate RP2 knockout iPSC using a simultaneous reprogramming and gene editing protocol (Howden et al., 2015). The location of the gRNA on exon 2 was selected due to its proximity to the c.358C>T p.R120X site, thus any knock out lines generated would closely mimic the consequences of this nonsense mutation. RP2 KO iPSC clones were identified through non-homologous end joining (NEHJ) mediated creation of indels; one clone with an 8bp deletion in exon 2 of *RP2* (RP2 c.371_378delAAGCTGGA; p.Lys124SerfsTer11) was selected for further study, as it had the shortest frameshift extension before a premature stop codon. Western blotting of iPSC showed efficient knockout, as no RP2 protein was detectable (Fig. 1 A, B).

**Figure 1.**
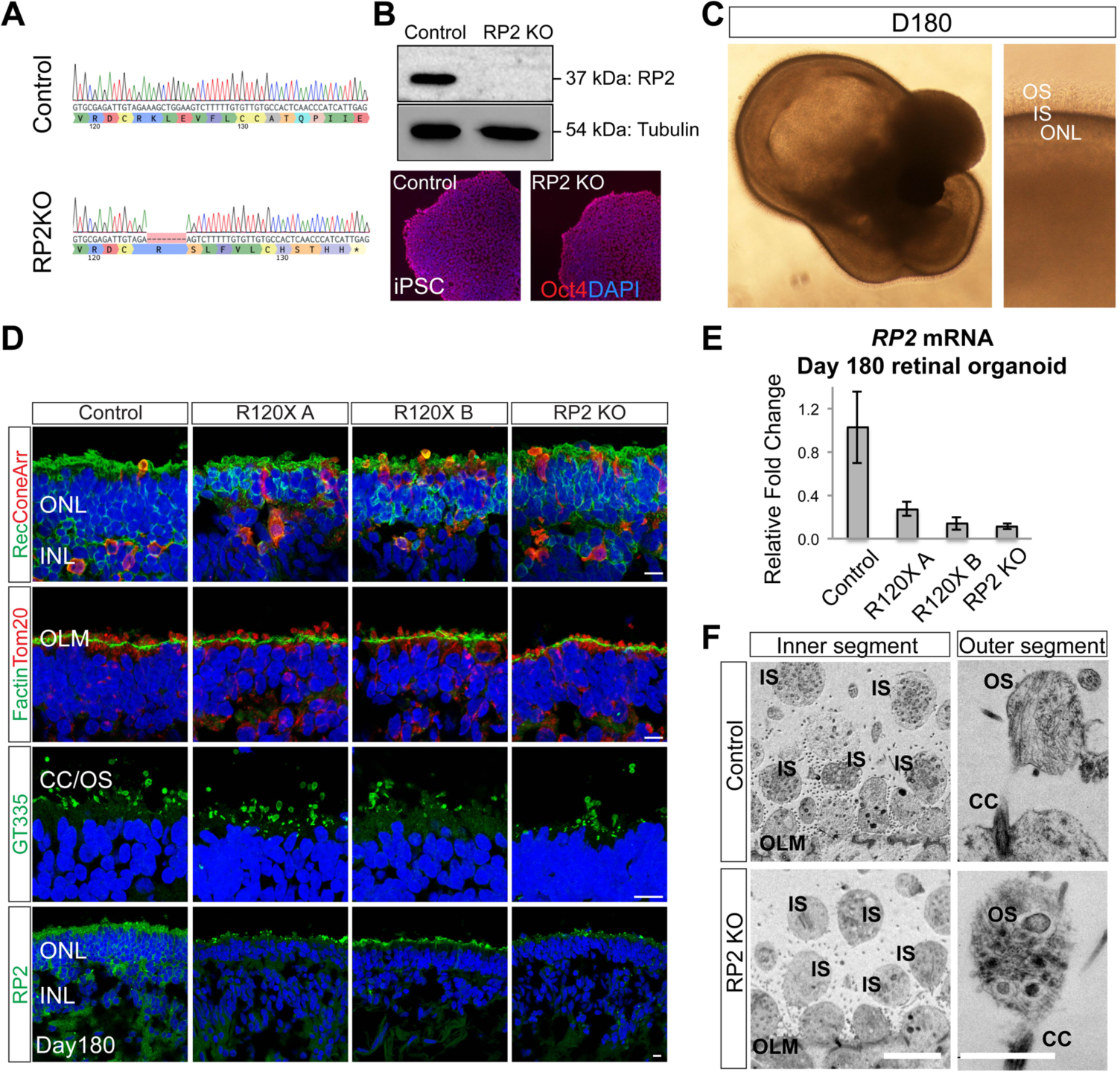
Retinal organoids from R120X, RP2 KO and isogenic control iPSCs. **A**. Sanger sequence trace of edited RP2 KO iPSC. CRISPR-Cas9 gene editing was used to create an 8bp deletion in exon 2 of *RP2* by NHEJ. **B**. Western blot and immunocytochemistry (ICC) of control and RP2 KO iPSC. **C**. Retinal organoid morphology at D180, a layer of IS/OS is visible by DIC above the transparent ONL. **D**. ICC of retinal organoids at D180. Photoreceptors (recoverin and coneArr), outer limiting membrane (OLM, F actin) mitochondria (Tom20) and connecting cilia/OS (GT335) expression in the ONL of control and RP2 null iPSC retinal organoids. RP2 is expressed at the plasma membrane of cells in the ONL and INL in controls organoids. Scale bar = 10µm **E**. qPCR of *RP2* mRNA in whole retinal organoids at D180 (n=3 independent organoids). Mean ± standard error of the mean (SEM). **F**. Electron micrographs of control and RP2 KO retinal organoids at D180.Scale = 5µm (left), 1µm (right).

The RP2 KO line, the non-edited isogenic control and two R120X patient iPSC lines were differentiated into retinal organoids (RO) over a period 4-10 months using previously described methods (Nakano et al., 2012, Zhong et al., 2014) with slight modifications. Both methods produced biologically similar organoids that were developmentally and structurally comparable at the timepoints tested. By Day 180 (D180) a transparent ONL with a brush like border was visible by light microscopy (Fig. 1C). Scattered rhodopsin positive cells were first detectable in control organoids in the recoverin positive ONL from D150 then increased in number over time as the ROs matured up to D180 (Fig. S1A). By D180 all cell lines were able to generate retinal organoids (ROs) consisting of a laminated structure with a compacted outer nuclear layer (ONL) containing recoverin and cone arrestin positive photoreceptors (Fig. 1D), above an inner nuclear layer (INL) containing PKCα positive bipolar cells and CRALBP/Nestin positive Müller glia (Fig. S1B). In all ROs, the ONL terminated at the apical edge, with an outer limiting membrane (OLM) that was strongly immunoreactive for F actin (Fig. 1D). Above the OLM, mitochondria (immunoreactive for TOM20) were enriched in globular inner segments (IS). Immunostaining for the ciliary marker ARL13B and polyglutamylated tubulin (GT335) revealed the bulging shape at the tip of the photoreceptor connecting cilia, as they matured to form outer segment (OS) like structures from D150 onwards (Fig. S1C). RP2 could be detected in control ROs at the plasma membrane in all cells in the inner and outer nuclear layers. In RP2 KO and R120X cell lines, RP2 immunoreactivity was absent by immunocytochemistry (ICC), although some background fluorescence was observed at the edges of the presumptive OS. qPCR was used to measure relative *RP2* mRNA expression in ROs at D180. The RP2 KO clone had only 20 % *RP2* mRNA relative to its isogenic control, similar to the R120X patient ROs (n=3) (Fig. 1E), suggesting that the mutant allele transcript is subject to nonsense mediated decay in all these ROs. Electron microscopy confirmed the presence of membranous rich structures at the apical ciliary tip, reminiscent of early OS formation, in both controls and RP2 KO ROs (Fig. 1F). These rudimentary OS were often found detached from the body of the RO indicating the flexibility, or fragility, of these structures in the absence of RPE. Collectively, these results show that RP2 ablation does not prevent the differentiation of iPSC into photoreceptors bearing OS-like structures in 3D RO culture.

### Loss of RP2 leads to photoreceptor cell death and ONL thinning

Similar to the neural retina *in vivo*, the outermost cell layer of the ROs consists of a uniform compacted ONL which terminates with photoreceptor synaptic pedicles that are separated from the inner retinal cells by a layer immunoreactive for synaptic structural protein Bassoon in the outer plexiform layer (OPL) (Fig. 2A). A reduction in the number of photoreceptor nuclei in the ONL, and thereby thickness, *in vivo* is a marker of photoreceptor cell degeneration. Therefore, ONL thickness was measured in isogenic controls and RP2 KO at D120, D150 and D180 (Fig. 2B). In control ROs, the average ONL thickness increased between D120 and D150 from 20 to 25 μm, then did not change significantly between D150 and D180. In contrast, in RP2 KO ROs the average ONL thickness decreased significantly between D150 and D180 (p=0.02). Similarly, R120X ROs from both patients had significantly thinner ONLs at D180 compared to the control cell line (p≤0.01; Fig. 2, Fig. S2). This suggested that photoreceptor cell death might be occurring between D150 and D180 in the RP2 null cell lines.

**Figure 2.**
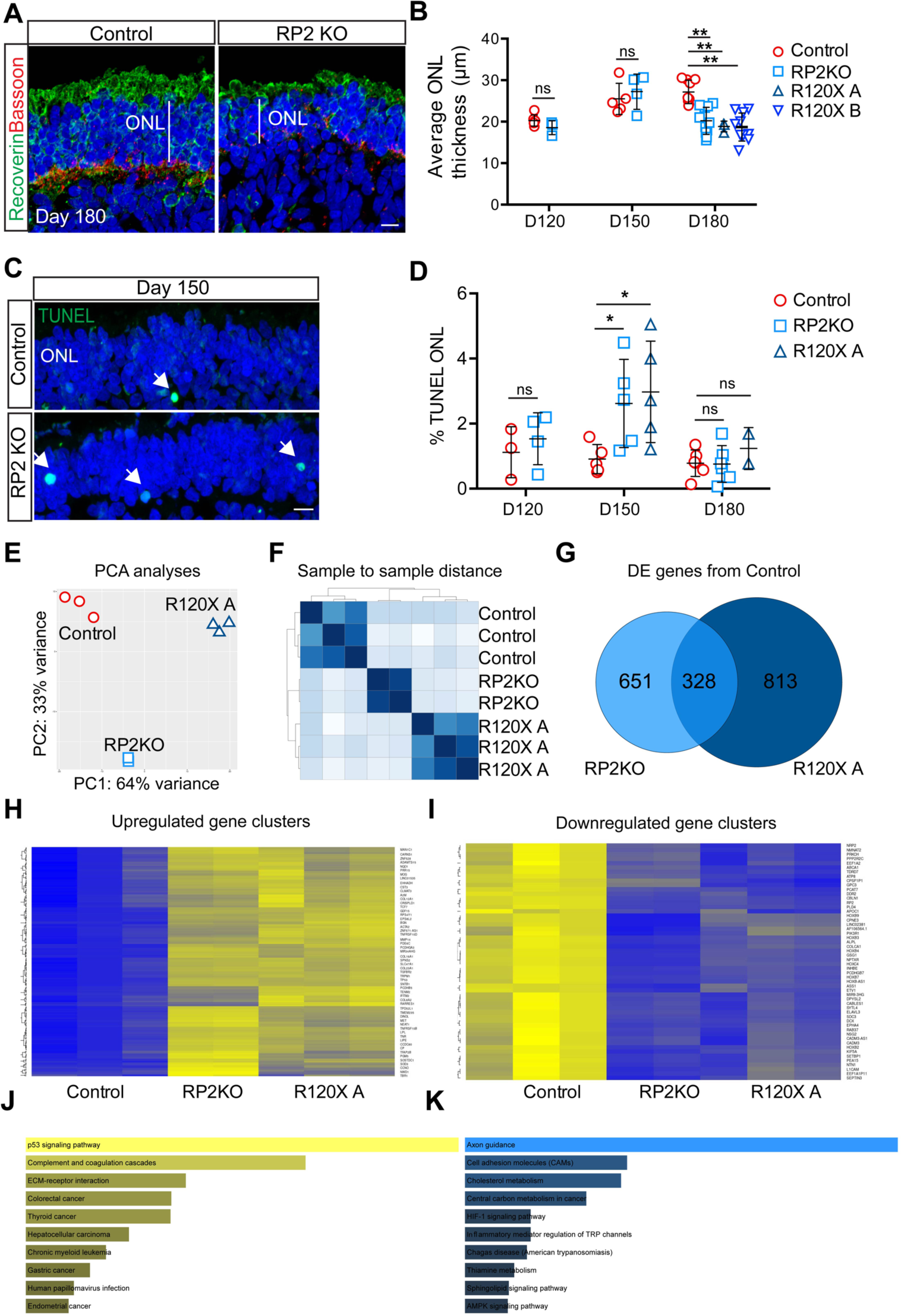
Photoreceptor differentiation associated cell death in RP2 KO organoids. **A**. ICC of control and RP2 KO retinal organoids showing reduced ONL thickness in RP2 KO. Recoverin staining demarcates the ONL terminating in the synaptic layer stained with Bassoon. Scale = 10μm. **B**. Mean ONL thickness per organoid was measured from tilescans of cryosections of a whole organoid at D120, D150 (n=5 control; n=4 RP2 KO at both time points), and D180. Significant ONL thinning was recorded at D180 in RP2 KO (n=9 independent organoids) and R120X lines (n= 3 R120X A organoids, n= 9 R120X B organoids), but not in controls (n= 10 independent organoids; p≤0.01; mean ± SD). **C**. TUNEL reactive nuclei (arrows) in the ONL of RP2 KO and isogenic control organoids at D150. Scale = 10μm. **D**. Quantification of TUNEL reactivity. RP2 KO organoids had a significantly higher proportion of TUNEL positive cells at D150 (n=5 independent organoids p≤0.05, mean ± SD) but not at D120 (n=3 control; n=4 RP2 KO) or D180 (n=5 control; n=6 RP2 KO). R120X organoids also had increased TUNEL reactivity at D150 (n=5 at D150 n=2 at D180 independent organoids). **E**. Principal component analyses of RNAseq data from ROs (n=3 control and R120X A, n= 2 RP2 KO independent organoids). **F**. Sample to sample distance between samples. **G**. Venn Diagram showing differentially expressed genes between RP2 KO and control and R120X and control and common genes. **H**. Heatmap showing upregulated clusters of DE genes (Blue lower expression, yellow higher expression). **I**. Heatmap of downregulated clusters of DE genes. **J**. KEGG pathway analyses of upregulated genes. **K** KEGG pathway analyses of down regulated genes.

To test this hypothesis further, we measured TUNEL reactivity across the ONL in RP2 KO and control ROs at D120, D150 and D180 (Fig. 2C, D). A small percentage of TUNEL positive nuclei were detectable in the photoreceptor ONL at all time points. In control ROs TUNEL reactivity was not significantly different at D120, D150 or D180. Whereas, the RP2 KO ROs had significantly higher percentage of TUNEL positive cells in the ONL at D150. There was no significant difference between RP2 KO and controls at D120 or D180, suggesting a peak of cell death during photoreceptor differentiation and maturation around D150 in RP2 deficient cell lines. This was confirmed in R120X-A ROs, with an increase in TUNEL reactivity at D150, which had resolved by D180 (Fig. 2D). These data show that there is a peak in photoreceptor cell death that correlates with their maturation and the time-course of increased rhodopsin expression.

### Gene expression changes associated with loss of RP2

To investigate gene expression changes that might be associated with the death of photoreceptors in the RP2 null organoids, RNAseq was performed on D150 controls, RP2-KO and R120X-A ROs. Principal component analyses (PCA) and sample to sample distance shows that the gene expression profile of the RP2 KO ROs were between the R120X-A patient line and the parental isogenic control (Fig. 2E, F). There were 328 shared differentially expressed (DE) genes between the RP2 KO and R120X ROs compared to control (Fig. 2G-I). By contrast there were 651 and 813 DE genes between control and RP2 KO and R120X-A ROs, respectively, that were not shared. KEGG pathway analyses of the shared DE genes revealed that the ‘p53 signalling pathway’ was the major upregulated pathway, whereas the major downregulated pathway was ‘axon guidance’ (Fig. 2J, K). KEGG pathway analyses of the axon guidance related changes showed that expression of genes in pathways that stimulate axon outgrowth, attraction and repulsion were reduced (Fig. S3). Whereas further investigation of the KEGG apoptosis and p53 signalling pathways showed that a number of pro-apoptotic genes, such as p21, BAX and PUMA were upregulated in both RP2 null ROs (Fig. S3), supporting the observation that loss of RP2 induces cell death in ROs.

### Rod photoreceptor differentiation and survival is compromised in RP2 KO and R120X retinal organoids

To identify which types of photoreceptor cells were most affected by the loss of RP2, rods and cones were stained with rhodopsin and cone arrestin at D180 (Fig. 3A). Strikingly, in the RP2 KO ROs there was reduced immunoreactivity for rhodopsin compared to isogenic controls. Unlike recoverin expression, which was widespread in the RP2 KO ROs, rhodopsin was restricted to patches of the ONL. To assess photoreceptor gene expression, we quantified rhodopsin (*RHO*) and rod transducin (*GNAT1*) expression by qPCR. In the controls both *RHO* and *GNAT1* showed an age-dependent increase, with a major increase between D150 and D180. In contrast, the RP2 KO ROs had reduced expression of *RHO* and *GNAT1* mRNA at D180, but not D120 or D150, confirming that this difference manifested from D150 onwards. The percentage of photoreceptors that were immunoreactive for rhodopsin were quantified as a percentage of DAPI positive nuclei across the full length of the ONL in sections from all ROs (Fig. 3C). The controls showed an increase in rhodopsin positive cells as the ROs matured. Despite variation between individual ROs, there was a significant difference in the percentage of rhodopsin expressing cells in mature ROs (D180) between the isogenic control and RP2 KO cell lines (Fig. 3A, C). Furthermore, the two R120X lines also showed very few rhodopsin positive photoreceptors at D180, similar to the RP2 KO ROs (Fig. 3C). By contrast, the percentage of cone arrestin positive cells was significantly increased in the RP2 KO and R120X ROs compared to controls at D180, and the mRNA (*ARR3*) was also increased in the RP2 KO ROs at D150 and D180 (Fig. S4), suggesting the defect primarily affects rods and not cones. There was no significant difference in bipolar cell numbers (Chx10/PKCα) between control and RP2 KO ROs (Fig. S4). In order to exclude that these differences might be attributed to simply a delay in the rate of maturation between the cell lines, control and RP2 KO retinal organoids were maintained in culture for a further 120 days to D300. The RP2 KO had fewer rhodopsin positive cells relative to the control ROs at D300, whereas recoverin positive photoreceptors were maintained (Fig. 3E).

**Figure 3.**
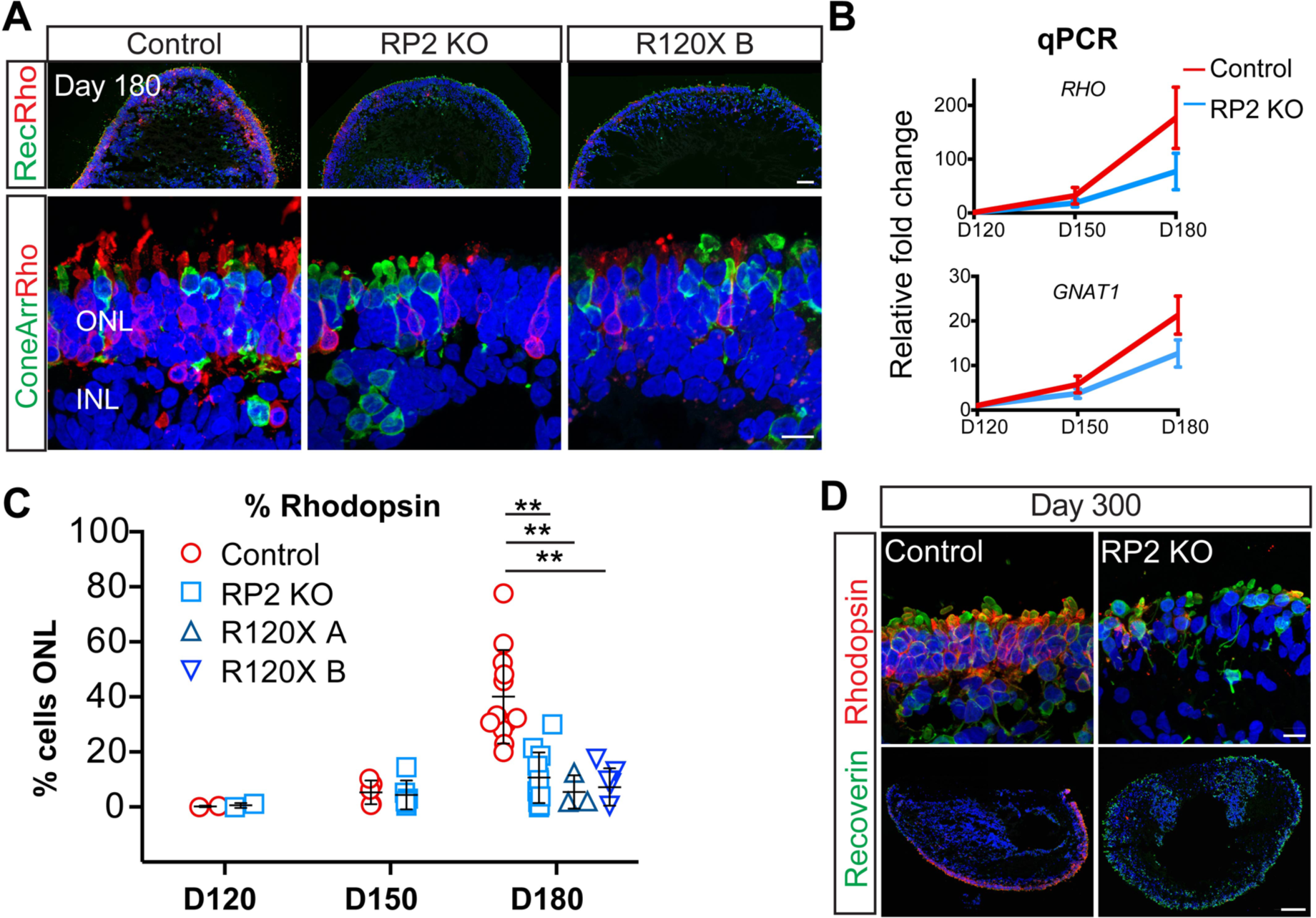
Reduced number of rod cells in RP2 KO and R120X patient retinal organoids. **A**. ICC of retinal organoids. Low (upper panel) and high (lower panel) magnification of recoverin, rhodopsin and cone arrestin immunoreactivity in the ONL at D180 in control, RP2 KO and R120X RP2 retinal organoids. Scale bars =50 μm and 10μm **B**. *RHO* and *GNAT1* levels in retinal organoids. qPCR showing relative fold change in mRNA in control and RP2 KO retinal organoids at D120, D150 and D180 (n=3,3,4 independent organoids). Mean ± SEM. **C**. Quantification of rhodopsin positive cells as a percentage of ONL in control, RP2 KO and R120X patient cell lines from D120-D180 of differentiation. Each data point represents the mean of 1 independent organoid, counts are from tilescans of whole organoid cross-sections (D120 n=2 control, n=2 RP2 KO; D150 n=5 control, n=6 RP2 KO; D180 n=12 control, n=11 RP2 KO, n=3 R120X A, n=5 R120X B; ** p≤0.01; mean ± SD). **D**. High and low magnification of control and RP2 KO organoids at D300 of differentiation stained with recoverin and rhodopsin (scale bar =10 μm upper panel, 100 μm lower panel).

### AAV2/5 efficiently transduces retinal organoids to augment RP2 expression in RP2 null photoreceptors

AAVs are able to transduce post-mitotic rods and cones in mice following subretinal injection (Sarra et al., 2002) and in iPSC derived rod and cone photoreceptors in ROs with varying efficiency (Gonzalez-Cordero et al., 2018). In order to assess the ability of AAVs to deliver RP2 to deficient photoreceptors, RP2 KO ROs were transduced with AAV2/5.CAGp.RP2 (Fig. 4A) at D140, before the observed onset of ONL thinning, and harvested at D180. At D180, RP2 protein could be detected by ICC throughout the ONL (Fig. 4B). At higher magnification, the RP2 signal was detected on the plasma membrane of photoreceptors (Fig. 4C). The percentage of RP2 positive cells was determined by scoring nuclei with closely adjacent RP2 signal on the plasma membrane that matched that cell morphology. The transduction efficiency was 90 ± 7% in the ONL (Fig. 4D). This included both rhodopsin positive rod cells (Fig. 4E) and cone arrestin positive cone cells (Fig. 4E). Sporadic RP2 staining in the inner retinal layers was also visible, suggesting a small proportion of the AAV is able to penetrate the RO and transduce inner retinal cells (Fig. S5). Analysis of mRNA levels by qPCR revealed an average 55-fold increase in RP2 transcript relative to control organoids (Fig. 4F).

**Figure 4.**
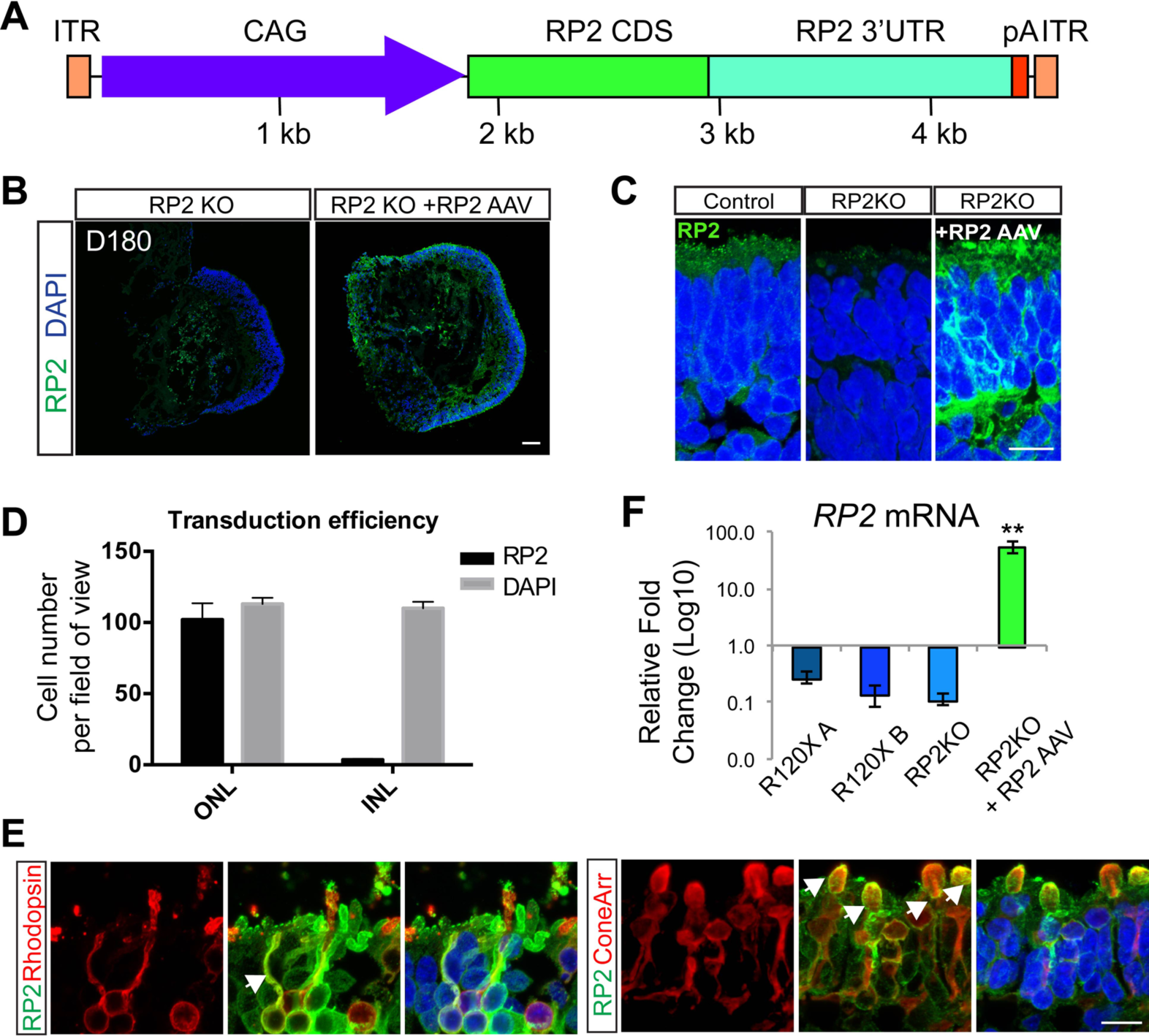
AAV2/5 RP2 efficiently transduces rod and cone photoreceptors. **A**. Schematic of AAV construction. CAG promoter (blue); RP2 CDS (green); RP2 3’UTR (teal); pA (red, minimal rabbit b globin polyA); ITR (orange). **B**. RP2 expression in RP2 KO retinal organoids following transduction. ICC on RP2 KO retinal organoid cryosections 6 weeks post transduction with AAV2/5 CAG RP2 showing RP2 expression across the photoreceptor layer (ONL). Scale bar = 50 μm. **C**. High power magnification of RP2 immunoreactivity. Scale bar = 10 μm. **D**. Cells with RP2 immunoreactivity in the ONL and INL scored against DAPI (mean = 90±7% ONL vs 3±0.4% INL, n=3 independent organoids; mean ± SD). **E**. ICC co-staining RP2 with cone arrestin or rhodopsin showing AAV driven RP2 expression in both rod and cone photoreceptors. Scale bar = 10 μm. **F**. qPCR of RP2 mRNA transcript levels in AAV transduced RP2 KO organoids relative to endogenous expression in control organoids (n=3 independent organoids; mean = 55 fold ± 7.7 SEM).

### AAV2/5 driven RP2 gene augmentation improves photoreceptor survival

To investigate if the increase in RP2 levels was altering the degenerative phenotype of the ROs, ONL thickness and rhodopsin immunoreactivity was compared in RP2 AAV transduced vs. untransduced RP2 KO ROs at D180 (Fig. 5A, C). The ONL was stained with recoverin and bassoon (Fig. 5A). Measurement of the ONL revealed that AAV-RP2 transduced RP2 KO organoids had a significantly thicker ONL than non-transduced controls at D180 (p≤0.01, Fig. 5B) with increased numbers of photoreceptors (Fig. S5). The ONL thickness post-transduction was similar to that observed for the isogenic control cells at D180, suggesting near complete rescue. In most, but not all, transduced organoids the percentage of rhodopsin positive cells was above the average for non-transduced RP2 KOs suggesting that AAV RP2 expression could restore rhodopsin immunoreactivity (Fig. 5D). This was also observed at the mRNA level, with up to a 3-fold increase in RHO level in transduced ROs; however, one transduced RO showed no response at the level of rhodopsin expression (Fig. 5E). The percentage of cone arrestin positive cells was also reduced following AAV transduction, but *ARR3* mRNA was unchanged (Fig. S5).

**Figure 5.**
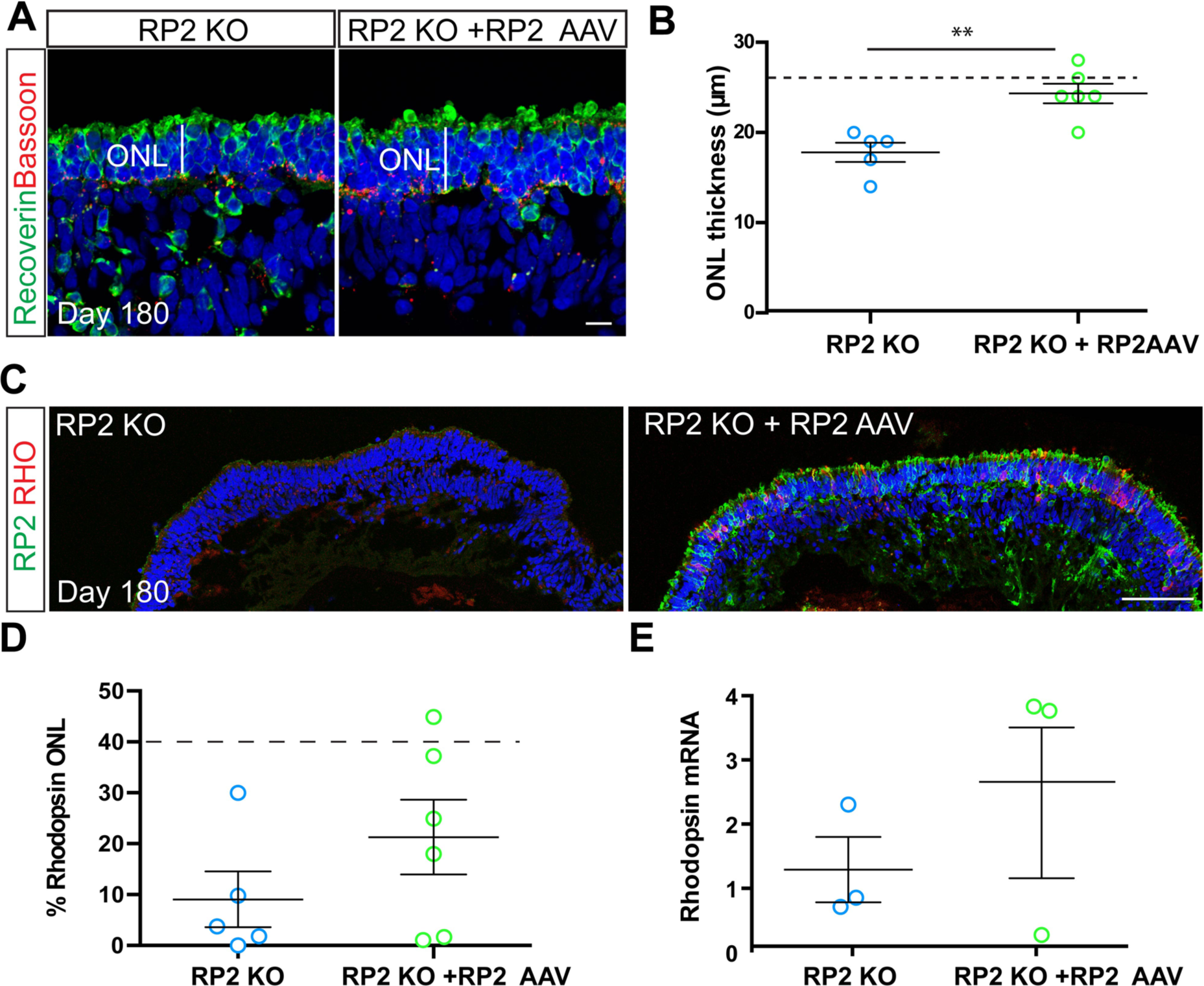
AAV2/5 driven RP2 overexpression rescues photoreceptor survival. **A**. ICC of retinal organoid ONL with recoverin (green) and bassoon (red) in RP2 KO control and AAV RP2 treated retinal organoids showing improved ONL thickness in AAV treated group. Scale bar = 10 μm. **B**. Quantification of ONL thickness in control and AAV transduced retinal organoids (n=5 RP2-KO, 6 RP2 KO+AAV independent organoids, ** p=0.016, mean ± SEM). Dotted line represents mean thickness in isogenic control parent cell line. **C**. ICC showing RP2 (green) and rhodopsin (red) expression following transduction. Scale bar = 50 μm. **D**. Quantification of ICC to assess rhodopsin positive in control and RP2 AAV transduced RP2 KO retinal organoids. Dotted line represents average control values (n=5 RP2 KO, n=6 RP2 KO + RP2 AAV independent organoids, p=0.23; mean ± SEM). **E**. qPCR for *RHO* mRNA levels in control and RP2 AAV treated RP2 KO retinal organoids (n=3 independent organoids). Mean ± SEM.

## Discussion

Here, we describe an *in vitro* model of RP2 XLRP and show that AAV gene augmentation of RP2 can successfully reverse measurable and clinically relevant disease phenotypes. The combination of iPSC reprogramming, CRISPR gene editing technology and AAV gene delivery has enabled a side by side comparison of RP2 KO, isogenic controls and XLRP patient ROs, to obviate some of the inherent variation in iPSC derived organoid models.

Differentiation of iPSC present an opportunity to probe genetic disease mechanisms in the cells and tissues which are affected by specific genetic changes. This is particularly useful in the case of ubiquitously expressed genes such as *RP2* that have disease pathology restricted to a specific tissue such as the retina. Here, we used iPSC reprogramming technology in combination with CRISPR/Cas9 gene editing to probe disease mechanisms in *RP2* XLRP.

All of the RP2 deficient cell lines used in this study successfully developed 3D ROs with advanced morphological features including lamination and formation of rod and cone photoreceptors. Strikingly, we observed a significant decline in photoreceptor cell numbers, specifically rhodopsin positive rods, and significantly thinner ONL after 150 days of differentiation in all RP2 deficient lines. Widespread rhodopsin expression is one of the later events in RO development, in keeping with the developmental time line *in vivo*; however, the presence of other late emerging cell markers in the RP2 KO and patient cell lines such as cone arrestin positive cone photoreceptors, PKCα positive bipolar cells, OS structures, together with the persistence of this rhodopsin deficient phenotype up to 300 days of differentiation suggests that this is not merely a case of developmental delay.

This phenotype has not been reported in any existing animal models of RP2 XLRP (Li et al., 2013, Zhang et al., 2015), suggesting that this is either specific to human retina, or to the *in vitro* organoid system. Despite the relative stability of ROs in comparison to *ex-vivo* retinal explant culture, which can be maintained for around 14 days (Johnson and Martin, 2008), it is likely that the conditions for retinal cell maintenance in culture are suboptimal. The absence of an opposing RPE monolayer, nutrient and oxygen deprivation through the lack of vascular blood supply, or the lack of connectivity for the inner retinal neurons, for example, may be sources of stress which accelerate the onset of disease phenotypes. This may go some way to explain the early ‘age’ of detectable changes relative to mouse and human RP2 XLRP *in vivo*.

In RP2 KO mice, ONL thinning is detectable only at 5 months of age in one model (Li et al., 2013), whereas almost no ONL thinning was observed at 12 months of age in another model (Zhang et al., 2015). Here we detected measurable differences in photoreceptor survival at the onset of rod differentiation and rhodopsin expression. Cell death is an important, non-pathological process during retinal development (Vecino et al., 2004), but is increased during differentiation in the photoreceptor cell layer of RP2 deficient ROs, linking rod maturation temporally with rod cell death in this model. Interestingly, ROs from an individual with a mutation in *RPGR*, the other major form of XLRP, also showed signs of photoreceptor cell death at around 150 days of RO differentiation, which was rescued by CRISPR mediated repair of the RPGR mutation in the iPSC (Deng et al., 2018). The induction of p53 pathways was also observed in the RPGR retinal organoid model (Deng et al., 2018), suggesting this could be a common pathway in both forms of XLRP in human ROs. Therefore, photoreceptor cell death appears to be a feature in ROs for some forms of IRD, but the precise mechanisms and whether there is an opportunity to inhibit the cell death process will require further investigation.

AAV is a highly efficient method of delivering therapeutic genes to photoreceptor cells and AAV2/5 has been shown to effectively transduce photoreceptors in a variety of species *inter alia* mice (Palfi et al., 2012), non-human primates (Boye et al., 2012) and human retinal explants (Wiley et al., 2018). The use of AAV in human ROs is currently limited, but studies have reported varying efficiency of transduction, which could be attributed to age of treatment, time of harvesting, vector tropisms, viral titre, the promotor used and sensitivity of detection of the transgene, e.g. antibody versus intrinsic fluorescence of a reporter (Gonzalez-Cordero et al., 2018, Quinn et al., 2019, Garita-Hernandez et al., 2020). Here we demonstrate highly efficient transduction using an AAV2/5 vector with RP2 under the control of a CAG promoter. The AAV appears to deliver RP2 more or less exclusively to the photoreceptor cells despite the use of a ubiquitous promotor. This may be due to a combination of vector tropisms and the organization of the organoid, where the photoreceptor layer is outermost and, therefore, in contact the viral particles in solution. Additionally, the OLM may prevent diffusion of AAV into the inner retinal layers. The interaction of AAV and photoreceptors in this system is analogous to the situation following subretinal delivery. Indeed, in a human treatment context, the optimal route of administration for this vector would be subretinal injection. Although intravitreal injection is less invasive, most AAV serotypes cannot transduce photoreceptors efficiently when administered intravitreally.

AAV RP2 was delivered at D140, prior to the onset of the increase in photoreceptor TUNEL reactivity and ONL thinning, but late enough to ensure efficient uptake, following reports of inefficient transduction at earlier time points (Gonzalez-Cordero et al., 2018). Not only was RP2 efficiently expressed at the mRNA and protein level, but it was also able to rescue the ONL thinning phenotype in RP2 KO retinal organoids implying a protective effect of RP2 overexpression in photoreceptor cells. Interestingly, although RP2 was expressed at over 55-fold the endogenous level, we did not observe any overt deleterious effects. In a previous study, some toxicity was observed with RP2 over-expression in mice 3 months post administration of the highest dose of AAV-RP2 vector (1e9 viral genomes (vg)/eye), although the level of RP2 over-expression in transduced mouse eyes was not defined (Mookherjee et al., 2015). As RP2 is a GAP for ARL3, it is possible that increasing RP2 to a high level would inhibit the production of ARL3-GTP and mimic the effect of loss of the ARL3 GEF, ARL13b, which causes Joubert syndrome; however, this might not occur in the cellular context because of the spatial separation of the proteins, with RP2 predominantly at the base of the cilium and ARL13b in the axoneme, which leads to a gradient of ARL3-GTP in the cilium. Despite very high levels of RP2, once ARL3 is trafficked into the cilium it can be converted to ARL3-GTP by ARL13b; however, this would not exclude other activities of over-expressed RP2 from disrupting homeostasis. Therefore, additional studies evaluating the therapeutic index for AAV-RP2 therapies in ROs, and/or the primate eye, would be of value, before translation to the clinic.

Although RP2 is expressed in multiple retinal cell types, it remains to be established whether optimal rescue of the RP2 disease phenotype may require expression of a replacement *RP2* gene in other retinal cell types, such as the RPE. The use of a ubiquitous promoter in the current study should, in principle, enable expression of the therapeutic gene in multiple cell types. Additional studies will be required to fully elucidate the requirement, or otherwise, for RP2 in the RPE, to develop optimal RP2 gene therapies. In contrast to the current study, Mookherjee et al. (2015) employed the photoreceptor-specific rhodopsin kinase promoter to drive expression of the RP2 gene, and achieved a partial rescue of the cone phenotype in a null RP2 mouse model but had no beneficial effect on the rod phenotype.

This study provides insights into the use of ROs for retinal disease modelling, with a phenotype related to loss of RP2 that is unique to the human retina cell culture model. Importantly, the photoreceptor cell death in the RP2 deficient organoids correlates with the timing of rod cell maturation and rhodopsin expression. Furthermore, we highlight how the use of AAV for the restoration of RP2 expression can reverse this phenotype, and as such could be further investigated as a potential therapeutic avenue for the treatment of XLRP.

## Author contributions

AL, KJ, CS, DO, ABP, NS, RG, MJH, NS, AP, NC performed experiments and/or analysed data; AL, KJ, CS, DO, NS, GJF, AJH and MEC conceived the hypothesis and designed experiments. AL, KJ, CS, GJF, AJH and MEC drafted the manuscript. All authors edited the draft manuscript.

## Acknowledgements

We are grateful to Robert Molday (UBC), Cheryl Craft (USC), Peter MacLeish (Morehouse School of Medicine) and Wolfgang Baehr (University of Utah) for providing antibodies. This work was funded by Moorfields Eye Charity through a generous donation (AJH, MEC), Fight for Sight (MEC, AJH), the NC3Rs (MEC) and the Wellcome Trust (MEC). It was also supported by the National Institute for Health Research Biomedical Research Centre at Moorfields Eye Hospital NHS Foundation Trust and UCL Institute of Ophthalmology (AJH is NIHR BRC Faculty). Support was also provided by the Health Research Board of Ireland (HRB), Fighting Blindness Ireland (FBI), the Medical Research Charities Group (MRCG) and Science Foundation Ireland (GJF).

## Experimental procedures

### Reprogramming and gene editing

iPSC were generated from two unrelated R120X individuals and control fibroblasts (BJ fibroblast ATCC® CRL-2522™) as previously described (Schwarz et al., 2015). RP2 KO iPSCs were produced by simultaneous reprogramming and gene editing using a method described previously (Howden et al., 2015). Guide RNAs were designed to target exon 2 of *RP2* (see supplementary data Table S1 for sequences), and were cloned into the pSpCas9(BB)-2A-Puro (PX459) V2.0 plasmid (Addgene plasmid 62988) according to a previously described protocol (Ran et al., 2013). iPSC clones were manually isolated, genomic DNA extracted (Promega) and PCR amplified with primers designed around the target site (Table S1). iPSC clones were analysed by Sanger sequencing to confirm *RP2* gene disruption. Off-spotter (https://cm.jefferson.edu/Off-Spotter/) was used to predict off-targets for the selected gRNA and the top 10 off-targets were assessed with Sanger sequencing, which detected no changes (Fig. S6A, Table S1). Additionally, all DE genes from the RNAseq analysis were cross-referenced with off-target predictions and a further 8 potential off-target sites with 4 or 5 mismatches were analysed (Fig. S6B, Table S1). These showed no sequence changes.

### Differentiation of iPSC to retinal organoids

RO differentiation was carried out as described previously (Nakano et al., 2012, Zhong et al., 2014). Briefly, iPSC were grown to near confluence in E8 media before detaching colonies in gentle dissociation buffer (StemCell Technologies) to form embryoid bodies (EBs). EBs were transitioned to neural induction media (NIM) in the presence of blebbistatin, before plating down at a density of approximately 20 EBs per cm2. Emerging transparent pouches of neuroepithelium were isolated using a needle and cultured in suspension in retinal maturation media + 0.5 μm retinoic acid up to D140 after which retinoic acid was removed. Alternatively EBs were generated by single cell dissociation and forced aggregation in 96 well V bottomed plates and cultured in suspension thereafter (Nakano et al., 2012). Days of culture are ± 3days.

### AAV production

The RP2 replacement vector comprising a CAG promoter, human RP2 CDS, RP2 3’UTR including a RP2 poly(A) sequence, and a minimal rabbit β-globin poly(A) was synthesized by GeneArt (Life Technologies) and cloned into pAAV-MCS using flanking NotI sites. More detail of sequences is in the supplemental information. Recombinant AAV2/5 viruses were generated by helper virus free, triple transfection (Xiao et al., 1998). Human embryonic kidney cells (accession number CRL-1573; ATCC, USA) were transfected with pAAV-RP2, pRep/Cap5 (Hildinger et al., 2001) and pHelper (Agilent Technologies, Inc., USA) at a ratio of 1:1:2, as previously described (O’Reilly et al., 2007). 72 hours post transfection, AAV particles were purified from the clarified lysate by differential precipitation with polyethylene glycol followed by caesium gradient centrifugation (Ayuso et al., 2010). AAV containing fractions were dialysed against PBS supplemented with Pluronic F68 (0.001%; Bennicelli et al., 2008). Genomic titres (viral genomes/ml; vg/ml) were determined by quantitative real-time PCR (qPCR; Rohr et al., 2002).

### AAV treatment

RP2 AAV was prepared to a final titre 4.73 E12 vg/ml. At 140 days, organoids with brush borders visible by light microscopy, which show inner and outer segment development, were transferred to a well of 96 well plate and incubated with 1E11 viral genomes in 75 μl media for 8 hours before topping the media up to 200 μl. Followed by 50/50 media changes every 2 days until Day 180.

### RNA extraction and qPCR

Retinal organoids or half retinal organoids were subjected to RNA extraction using RNeasy MicroKit (Qiagen) and cDNA synthesis was performed using Tetro cDNA synthesis kit (Bioline). qPCR was carried out on an Applied Biosystems 7900HT Fast Real-Time PCR system using the SYBR Green method using 1 μl cDNA per triplicate. Data from the qPCR was normalized to the geometric mean of the expression of two internal reference genes in each sample. *POLR2A* and *MAN1B1* were chosen due to their consistency across the sample groups, which was determined using the GeNorm algorithm (Vandesompele et al., 2002). Primers were designed to cross exon boundaries (See Table S2).

### RNAseq

Three ROs from BJ controls and R120X cell lines and two ROs from the RP2-KO cell line were harvested at D150. The RNA was extracted with the RNeasy-micro kit (Qiagen) following manufacturer instruction followed by paired-end sequencing at 100M read depth for each sample (Illumina, Otogenetics, Atlanta - GA, US). Raw .fastq sequences were cleaned from any residual sequencing adapter using cutadapt with parameters -m 20 and -e 0.1. Fragments were then aligned to the human genome (build 38, Ensembl version 92) using STAR and genes counted using featurecounts. RNAseq data is available GEO: GSE148300.

Differential expression (DE) analysis was then performed using the DESeq2 pipeline. Initial inspection of the dataset showed biases, which were blind estimated and corrected using the sva package (DESeq2 manual). The DE analysis showed 1141 genes were differentially expressed between the R120X retinal organoids and the control. Whereas 979 genes were differentially expressed between the RP2-KO ROs and the isogenic control and 328 genes were common to the two RP2-depleted backgrounds. Of these, 232 genes were consistently upregulated and 56 were downregulated. The list of DE genes is shown in Table S3. Finally, we used the web platform Enricher to source any significant pathway (https://amp.pharm.mssm.edu/Enrichr/). According to the Kyoto Encyclopedia of Genes and Genomes (KEGG) there was a significant upregulation of the ‘p53 signalling pathway’ in the RP2-KO and RP2-R120X retina (Fig. S3).

### Immunofluorescence

ROs were either fixed whole or bisected under a dissecting microscope using micro scissors (Fine science tools) with the other half processed for RNA extraction. Retinal organoids or half retinal organoids were fixed in 4% PFA at 4 °C for 20-30 minutes before cryoprotection by immersion overnight in 30% sucrose/PBS. Following orientation under a dissecting microscope in OCT they were frozen and cryosectioned into 6 μm sections. For ICC, slides were blocked in 10 % donkey serum, 0.1 % TritonX for 1 hr before incubation with primary antibodies for 2 hrs, (see Table S4 for primary antibodies) and donkey anti Rabbit or mouse AlexaFlour 488 or 555 secondary antibodies for 1 hr (Thermofisher). Nuclei were visualized using DAPI (2 μg/ml) staining for all images.

### Imaging

All images were obtained using Carl Zeiss LSM700 or LSM 710 laser-scanning confocal microscope. Images were exported from Zen 2009 software and prepared using Adobe Photoshop and Illustrator CS4. All measurements were performed in Fiji (Schindelin et al., 2012.) For quantification of TUNEL, rod and cone cell numbers, and ONL thickness measurements the tilescan function was used to obtain images of whole organoid sections at 40x magnification (more detail in supplemental information). Individual rod, cone and TUNEL positive cell numbers in the ONL were manually counted across the whole section. ONL thickness was calculated by measuring the length and area of the whole recoverin positive ONL in the DAPI channel using Fiji.

### Electron microscopy

Retinal organoids were fixed overnight in 3% glutaraldehyde and 1% paraformaldehyde buffered to pH 7.4 with 0.08 M sodium cacodylate-HCl buffer. After rinsing in 0.1 M sodium cacodylate-HCl buffer (pH 7.4) twice for 5 minutes, the retinal organoids were post-fixed in 1% aqueous osmium tetroxide for 2 hours, dehydrated by passage through ascending alcohols (10 minute steps, 1x 50-90% and 3 x 100%) and two changes of propylene oxide, and infiltrated overnight with 1:1 mixture of propylene oxide:araldite on a rotator. Finally, retinal organoids were infiltrated with araldite resin over 4-6 hours and embedded in fresh resin, which was then cured by overnight incubation at 60°C. Semithin sections (0.75μm) were stained with a 1% mixture of toluidine blue-borax in 50% ethanol, and ultrathin sections were contrasted with Reynolds lead citrate for imaging in a JEOL 1010 TEM operating at 80kV. Images were captured using a Gatan Orius CCD camera using Digital Micrograph software.

### Statistical analysis

Statistical analysis was carried out on GraphPad Prism. Data was subjected to 2-way ANOVA and/or by multiple T tests using the Holm-Sidak method, with alpha=5.00%. Each time point was analysed individually, without assuming a consistent standard deviation (SD). Significance was determined at a p value of less than 0.05.

