## Supplemental Information for "Modelling and rescue of RP2 Retinitis Pigmentosa using iPSC Derived Retinal Organoids"

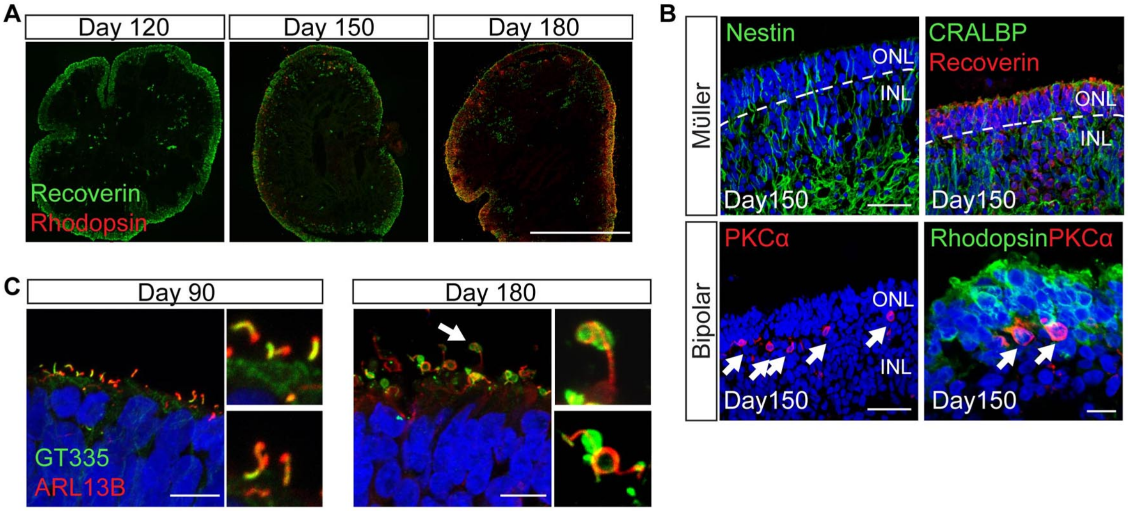


**Figure S1.** ***Control retinal organoid differentiation.*** Related to Figure 1.

**A.** ROs show increased rhodopsin expression between D150 and D180 Scale bar =500 μm.

**B.** ROs contain Muller glia, stained with nestin (green) and CRALBP (green) and bipolar cells, stained with PKCα Scale bars left = 50 μm; lower right = 10 μm**.**

**C.** Apical cilia in ROs stained with ARL13b (red) and GT335 (green) Scale bars = 10 μm.

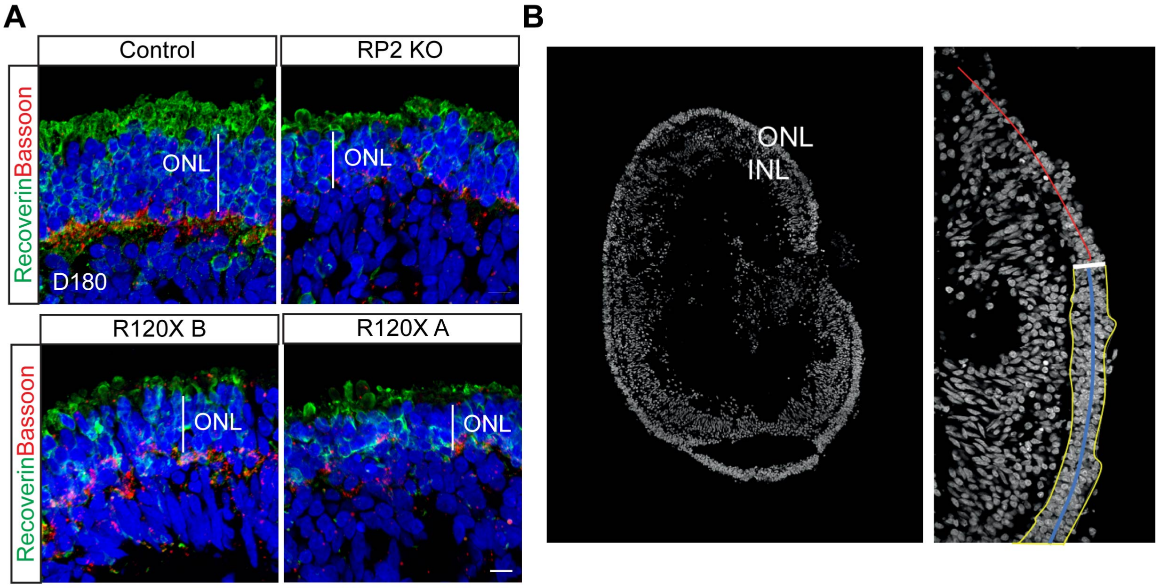


**Figure S2.** ***Thinning of the ONL in RP2 null cells.*** Related to Figure 2.

**A**. Representative images showing reduced ONL thickness in RP2 KO, R120X A and R120X B ROs. Scale bar = 10 µm

**B**. Tilescans of retinal organoid to illustrate the method of ONL measurement. Whole organoid (left panel) close up of ONL (right panel). The area of the ONL is enclosed within the yellow marked area and the length of ONL indicated by the blue line. The red line indicates an area of non-photoreceptors that were not included.

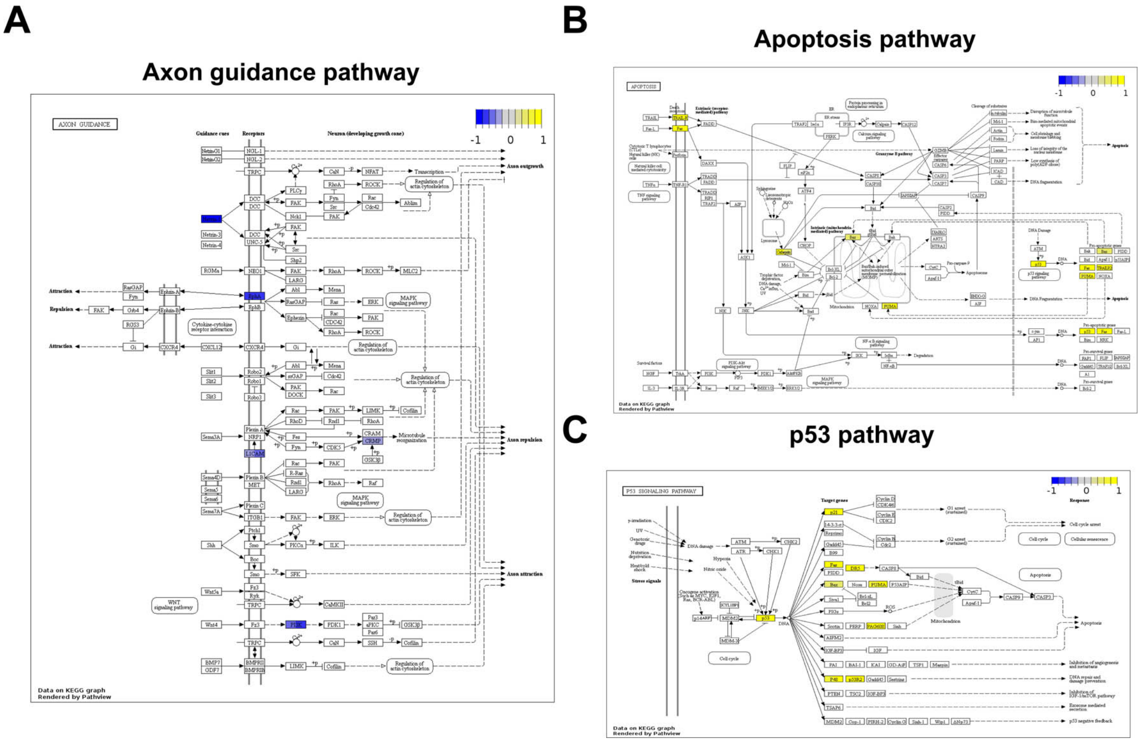


**Figure S3.** ***KEGG analyses of shared DE genes between RP2 nulls and control.*** Related to Figure 2.

**A.** KEGG analyses of RP2 KO and RP2 R120X common downregulated DE genes compared to control ROs at D150 in 'axon guidance' pathways.

**B**. KEGG analyses of RP2 KO and RP2 R120X common DE genes that are upregulated compared to control ROs at D150 in the 'apoptosis' pathway.

**C**. KEGG analyses of RP2 KO and RP2 R120X common DE genes that are upregulated compared to control ROs at D150 in the 'p53' pathway

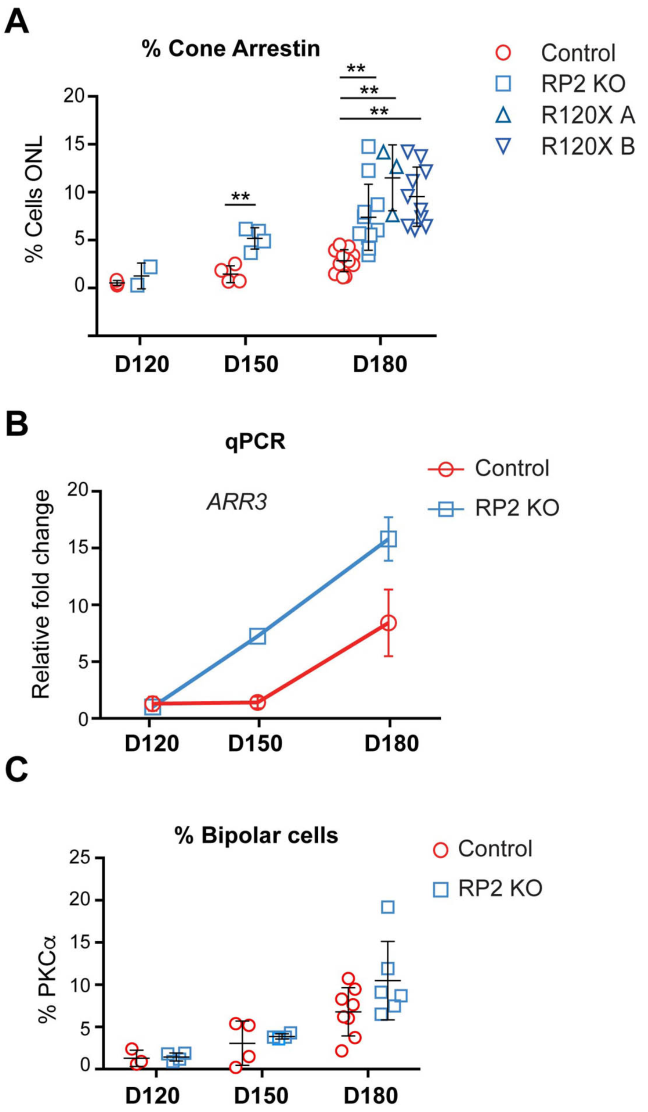


**Figure S4**. ***Cone photoreceptor and bipolar cells in RP2 null organoids.*** Related to Figure 3.

**A.** Percent of cells in ONL positive for cone arrestin immunostaining, Control (red circle; n=3, D120; n=4 D150 n=10 D180); RP2 KO (blue square; n=2, D120; n=4 D150 n=10 D180) R120X A (blue triangle; n=3 D180), R120X B (blue inverted triangle; n=10 D180); ** $p<0.01$ each n represents the mean from a full tilescan of an independent organoid. Mean ±SD

**B**. *ARR3* levels in retinal organoids. qPCR showing relative fold change in mRNA in control and RP2 KO retinal organoids at D120, D150 and D180 (n=3,3,4 independent organoids respectively; mean ±SEM).

**C**. Cell counts of bipolar cells (positive for PKCα) in control and RP2 KO ROs showed no significant differences in bipolar cell numbers at D120, D150 or D180 (controls n=3, 4 and 8 RP2 KO n=4, 4, 6, mean of tilescan of full independent organoids; mean ±SEM).

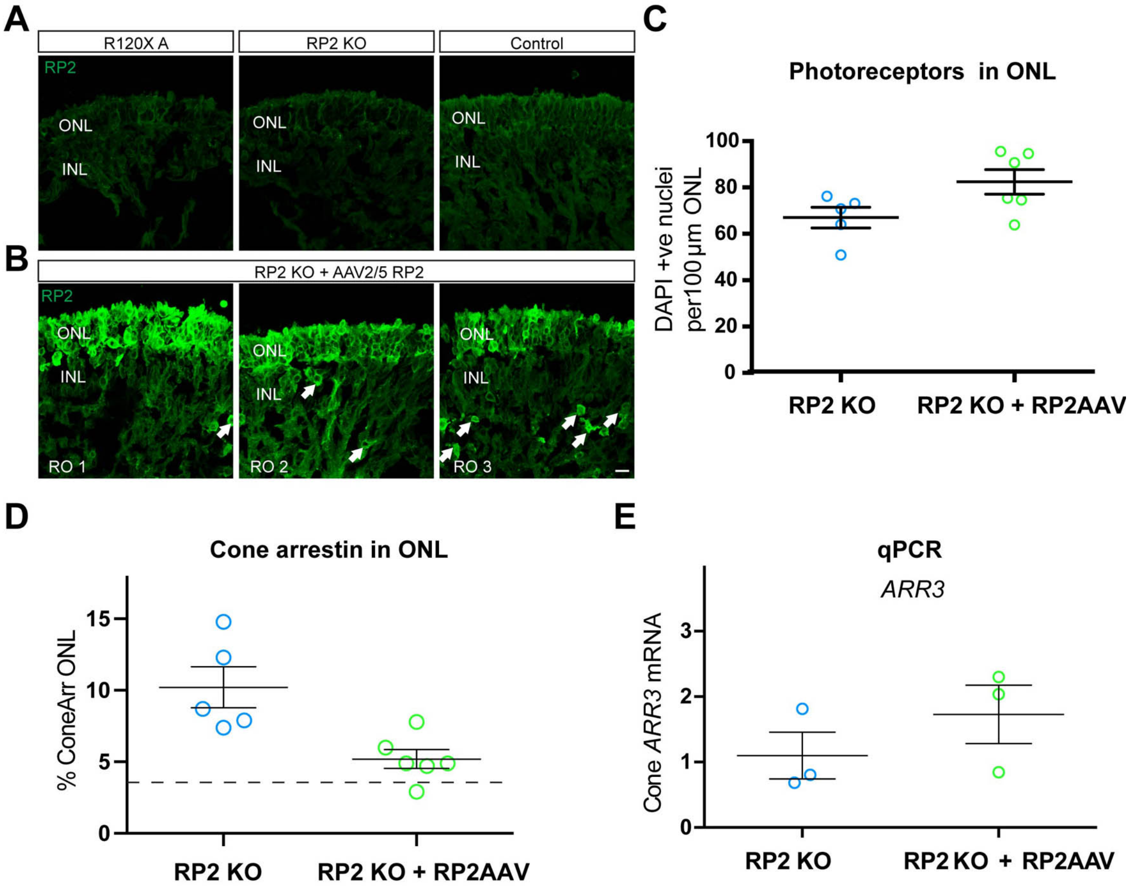


**Figure S5.** ***Transduction of RP2 KO organoids with RP2-AAV.*** Related to Figure 5.

**A**. ICC for RP2 in untreated D180 RP2 R120X A and RP2 KO and control ROs.

**B**. RP2 immunostaining after RP2-AAV transduction in 3 different organoids (RO1, RO2 and RO3). The few strongly positive cells for RP2 outside of the ONL are indicated with arrows. Scale bar = 10 µm.

**C**. The number of photoreceptors in the ONL. Mean of nuclei in the ONL per 100 µm (n=5 RP2 KO and n=6 RP2 KO + RP2AAV independent organoids for each condition; p=0.058; mean ±SEM).

**D**. Percentage of cells in the ONL that were positive for cone arrestin staining in untreated and after RP2-AAV transduction at D180, the dotted line indicates the mean percent in controls (RP2 KO n=5, RP2+RP2AAV n=6 independent organoids; p=0.055; mean ±SD).

**E**. *ARR3* levels in retinal organoids following RP2-AAV transduction. qPCR showing relative fold change in mRNA at D180 (n=3 independent organoids for both conditions; mean ±SEM).

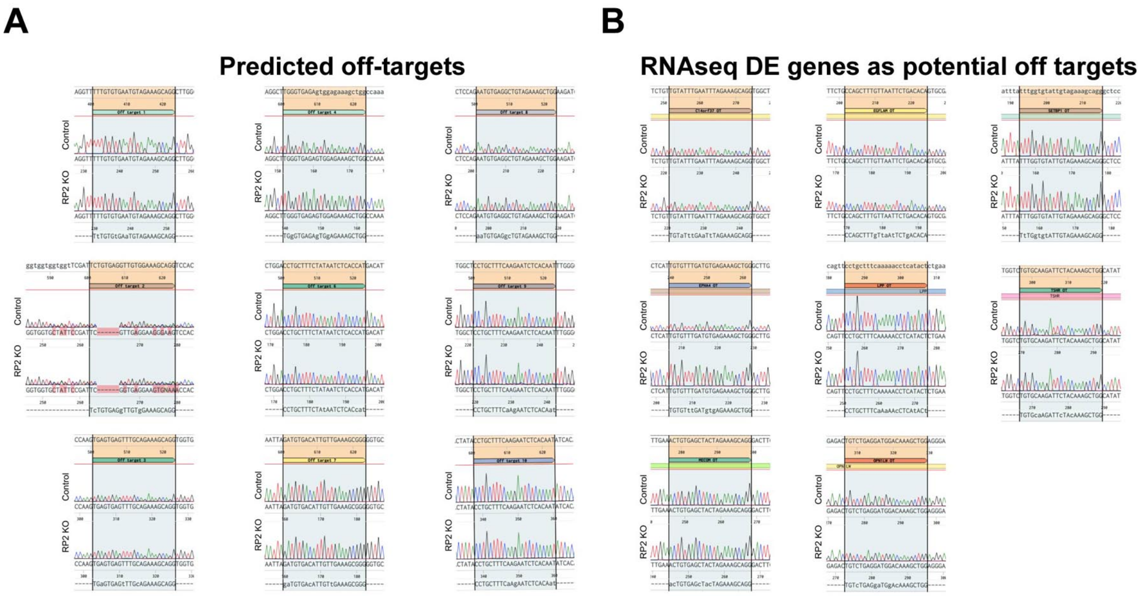


**Figure S6.** ***No off-target editing was observed in the RP2 KO.*** Related to Figure 1 and experimental methods.

**A.** Sanger sequencing results of Top 10 predicted off-targets for RP2 gRNA2. Possible off-targets for the RP2 gRNA2 were predicted using the Off-Spotter online tool (https://cm.jefferson.edu/Off-Spotter/). Sequences from the RP2 KO line and the isogenic control line were aligned using Benchling (www.benchling.com). No mutations were observed in any of the predicted off-target sites. Off target 2 happens to occur in a region where the control line has a naturally occurring variation. Primers for Off target 5 failed to amplify the DNA due to low complexity and high GC content in the target region. Lower case letters in the off-target sequences below the traces indicate the sites at which the off-targets vary from RP2 gRNA2.

**B**. Sequencing results of RNAseq DE genes with possible similarity to the gRNA sequence. The list of DE genes generated by RNAseq analysis of RP2 KO vs control ROs was cross referenced with the list of Off-targets for RP2 gRNA2 (Off-spotter online tool; 431 off-targets predicted). Sanger sequencing revealed that none of these sites showed any editing by off-target Cas9 activity.

**Table S1. gRNA, Primer and Off-target sequences.** Related to Figures 1, 4, and S6

| Target | Forward | Reverse |
|---|---|---|
| **gRNA sequence for RP2 exon 2 CRISPR** | | |
| *RP2* gRNA2 | TGTGTGAGATTGTAGAAAGC**TGG** | |
| **PCR primer sequences** | | |
| *RP2* exon2 | CCAAAAGACTACATGTTCAGTGGA | GCATGATTCATCGCTGCTCT |
| **Off-target PCR Primer sequences** | | |
| OT1 – Chr13 + 99786425 - 99786447 | GGTCAGGGCCAGCACATAAT | TAGCTAATGTGCCCTCAGCG |
| OT2 - Chr 6 + 38143360 – 38143382 (BTBD9 pre-mRNA) | ACCACGAGGGAGTTAGAGGA | AGCCACATGAGGTTGAAGTCC |
| OT3 - Chr 9 + 91974085 – 91974107 (SECISBP2 pre- mRNA) | AGTTCTTGCCTGTGTGTCTGT | TAGCAGAACCCCCACTACGC |
| OT4 - Chr 7 + 157998199 – 157998221 (PTPRN2 pre-mRNA) | TAGCCAAAGCTCTTGCTTCAGT | GTGCCCTATCCTACTTGGCTG |
| OT5 - Chr 3 – 132906887 -132906909 (TMEM108 pre-mRNA) | CTGTCAAGGAAGGAAGGGAGG | GGAGAAAGGGCCCCCTATAA |
| OT6 - Chr 5 – 116400519 - 116400541 | AACTGTGTCCCACAATAAGGGG | TGCTGGGACCTCCTCAAATTC |
| OT7 - Chr X + 119452379 - 119452401 | AAGCCCACAGGAACTGGGTT | AAACCAAAGTAGGTGGGTCCT |
| OT8 - Chr 10 + 22037281 - 22037303 | ATTCTGCCGGGGTTTAGAGC | AGCAAGGTGAACTGACACTGG |
| OT9 - Chr 1 – 23887266 - 23887288 | AGTAGCTCCAGCTCCATCTGA | ATCTTTCTTGGGCCACAGAGC |
| OT10 - Chr 11 – 109695393 - 109695415 | AAAATGGCTATGCAGCACAAGTT | TTAGCTTCTCCCTTACACCTGAAC |
| **RNAseq DE gene potential off-target PCR Primer sequences** | | |
| C14orf37 | ACTGGGCTTTGAAAGATGGAGA | AGGCACTCAAATGGTGGAGC |
| EGFLAM | TAGGGCTTTGCACATCGCA | CGGGTTTAGTGGGAAGGCAG |
| EPHA4 | CCTTGTCGATCTTCCACTCCC | ATGACGTGTGGAGCACTGAC |
| LPP | TCCGACTTTGCCTTCCTTCT | CTCTGTGGCACCTATCACAACT |
| MECOM | AGCTGCCATTGCTTTTTCTCA | TGCTTAGGAAACACACGCCT |
| OPN1LW | TGGGTAGTGGTCTTCCCCTC | AGCCCATTTGTGAAGACGGA |
| SETBP1 | TTTCATGCCAGTGACCTTCG | TTGCTCATTGACTTTTGACATGGG |
| TSHR | AGCCACAACCACTCTTGACT | TGACTGGTTTTCAGGTGCTAGAT |

**Table S2. Primer sequences for qPCR.** Related to Figures 3, 4, 5 and S4, and S5

| qPCR primer sequences | | |
|---|---|---|
| **Target** | **Forward** | **Reverse** |
| RHODOPSIN | GGTGGTGTGTAAGCCCATGA | CCTCGGGGATGTACCTGGAC |
| GNAT1 | CTGCTCACTCTGTCCCTTCG | TTGACGATGGTGCTCTTCCC |
| RP2 EX1-2 | GACCATGGGCTGCTTCTTCT | ACTGTTGTCCTGCTACCGTC |
| ARR3 | CATGTGCCTTTCGCTATGGC | TCAATCCCACAGGGCTTTCC |
| MAN1B1 | ACCGTGGAGAGCCTGTTCTA | GTTTGGGTCATCGGAGAAGA |
| GAPDH | CCCCACCACACTGAATCTCC | GGTACTTTATTGATGGTAC |
| POLR2A | GCGGGGTGAAGTGATGAAC | ATCATCGGGATGGGTGCTGT |

**Table S3. List of DE genes from D150 RNAseq analyses.** Related to Figure 2.
See excel file Supplementary Table S3.xls.

**Table S4. List of antibodies used.** Related to all figures.

| Marker | Species | Dilution | Supplier | Catalog number |
|---|---|---|---|---|
| Oct4 | Rabbit | 1:1000 | Abcam | ab19857 |
| Nanog | Mouse | 1:1000 | Thermo Invitrogen | MA-1-017 |
| RP2 (western blot) | Sheep | 1:2500 | Sheep serum | References 1, 2 |
| RP2 (ICC) | Rabbit | 1:200 | ProteinTech | 14151-1-AP |
| Recoverin | Rabbit | 1:500 | Millipore | AB5585 |
| Cone Arrestin (7G6) | Mouse | 1:100 | Gift from Wolfgang Baehr | |
| Rhodopsin (4D2) | Mouse | 1:500 | Millipore | MABN15 |
| F-actin | AlexaFluor 488 Phalloidin | | Thermo Invitrogen | A12379 |
| TOM20 | Mouse | 1:150 | Santa Cruz | SC17764 |
| GT335 | Mouse | 1:800 | Adipogen Life Sciences | AG-20B-0020 |
| Arl13B | Mouse | 1:1000 | Abcam | ab136648 |
| Bassoon | Mouse | 1:500 | Enzo | SAP7F407 |
| Nestin | Mouse | 1:200 | Abcam | ab22035 |
| CRALBP | Mouse | 1:300 | Thermo Invitrogen | MA1-813 |
| PKCα | Rabbit | 1:200 | Abcam | ab32376 |

### Supplemental Experimental Procedures

**AAV production**
The following sequences were used in the construct: CMV IE enhancer (661-1024 bp from X03922), Ch-B-Actin proximal promoter/intron 1 (251-1542 bp of X00182; 1538-39 bp AG was changed to CA); RP2 CDS (190-1270 bp of pNM_006915.2), RP2 3'UTR (1271-2440 bp of NM_006915.2), RP2 poly(A) (50341-50580 bp of NG_009107.1) and minimal rabbit B-globin poly(A) (Levitt et al., 1989)[3].

**ONL measurement methods**
Mean ONL measurements for control, RP2 KO and R120X patient derived ROs were assessed at three time points (D120, D150 and D180). Each data point represents the mean ONL measurement for an individual unique organoid. Mean ONL measurements were performed on confocal images taken with a 40x objective on a Zeiss LSM710. Organoids varied in size but approximately 12-20 images were acquired with the 40x objective so that the entire organoid section was imaged in a series of tiles. Each tile consisted of a maximum intensity projected Z stack. These tiles were then merged to produce a composite of the entire ONL in each organoid and processed in image J (FIJI). In mature ROs the ONL layer is easily determined by the organisation of nuclei in the DAPI channel, but this was also verified by examining the termination of photoreceptor marker staining (e.g. recoverin, rhodopsin, cone arrestin). Furthermore, between the densely packed nuclei of the ONL there is a physical gap to the INL (Fig. S2), this synaptic outer plexiform layer is labelled with bassoon in Fig 2A and S2. In imageJ (FIJI) the polygon selection tool was used to measure the area around the entire organised ONL in the DAPI channel (yellow line in Fig. S2B). In organoids where the full section was not organised into ONL/INL (e.g if an uncharacterised non-retinal area was present) the junction between retina and non-retina where the ONL tapers away was excluded in all cases (red line in Fig. S2B). In sections where 2 or more retinal 'lobes' were separated by non-retinal tissue, the lobes were measured separately and averaged into one number. The segmented line tool was used to draw a line through the centre of the selection area and the length (in pixels) measured (blue line Fig. S2B). The mean ONL thickness was then determined by division of the total area by the length. In this way deviations in ONL thickness throughout the organoid were accounted for. The final value (in pixels) was then converted to micrometers using for the scaling information from the acquisition format and 40x objective used.

**TUNEL assessment method**
The percentage of TUNEL positive nuclei in the outer nuclear layer for control, RP2KO and R120X patient derived organoids was derived over three time points, D120, D150 and D180 (Fig. 2D). Each data point represents TUNEL data from an individual unique organoid. TUNEL measurements were carried out on confocal images taken with a 40x objective on a Zeiss LSM710. Organoids varied in size but approximately 12-20 tiles at the 40x objective were acquired so that the entire organoid section was imaged. Each tile consisted of a maximum intensity projected Z stack. These tiles were then merged and opened in image J (FIJI). The area comprising ONL was determined as described above. DAPI nuclei numbers were counted manually in image J. Nuclei were counted in a representative area of ONL for each organoid and the number extrapolated based on total ONL area for each tile scan. TUNEL reactive nuclei within the predetermined ONL area were counted manually in the appropriate channel. The % TUNEL positive nuclei per organoid was then calculated. Across all data points the mean number of nuclei included per organoid was 1522 and the number of TUNEL positive nuclei counted was between 1 and 43.
